# Neurometabolic signatures of addiction vulnerability and heroin versus social seeking: a PET study in rats

**DOI:** 10.64898/2026.03.19.712973

**Authors:** Ginevra D’Ottavio, Alana Sullivan, Emma Pilz, Ingrid Schoenborn, Oscar Solis, Juan Gomez, Thorsten Kahnt, Michael Michaelides, Yavin Shaham

## Abstract

Only a subset of heroin users develop addiction, characterized by binge-like heroin use and preference for heroin over other rewards, including social rewards. We recently established a rat model of these features.

We trained rats to lever-press for social interaction and heroin (or saline, control) infusions and then tested heroin- and social-seeking and heroin-vs.-social choice. During 3-5 abstinence weeks, we used 2-deoxy-2-[¹⁸F]fluoro-D-glucose (FDG) PET imaging to assess regional brain metabolic activity at rest (homecage) and during heroin and social seeking. We assessed regional differences in FDG uptake using unbiased voxel-wise analysis and statistical parametric mapping, and correlated FDG uptake with principle-component-analysis-derived addiction severity score incorporating heroin intake, binge-like episodes, and heroin preference.

Compared with saline-trained rats, heroin-trained rats showed overall higher FDG uptake across multiple brain regions at rest and during both reward-seeking tests. Comparison of heroin-vs.-social-seeking in heroin-trained rats showed higher uptake in claustrum/lateral striatum and auditory cortex during social seeking. Analysis of individual differences showed that addiction severity was primarily associated with metabolic alterations under resting conditions rather than during heroin- or social-seeking. At rest, higher addiction severity was associated with lower uptake in piriform cortex and higher uptake in ventral hippocampus, whereas during heroin-seeking, addiction severity was associated with lower uptake in post-subiculum and cerebellum. Addiction severity was not associated with differences in social seeking or FDG uptake during social seeking.

These findings identify neurometabolic features of social and heroin seeking and heroin addiction vulnerability that can potentially serve as brain biomarkers and targets for neuromodulation.

**Significance Statement:** Heroin addiction develops in only a subset of users, yet the determinants of vulnerability versus resilience to addiction remain largely unknown. We combined a rat model capturing key features of heroin addiction, including binge-like heroin intake and preference for heroin over social interaction, with behavioral heroin- and social-seeking assays and longitudinal whole-brain metabolic imaging using FDG-PET. We identified distinct patterns of neurometabolic alterations associated with heroin self-administration and addiction severity at rest and in the context of heroin seeking. In contrast, heroin self-administration and addiction severity were not significantly associated with neurometabolic alterations during social seeking. These findings highlight brain-wide neurometabolic features of vulnerability to heroin addiction that can serve as brain biomarkers and targets for neuromodulation.

## Introduction

Opioids are among the most effective analgesics for pain management worldwide (1). Although medical exposure alone rarely leads to iatrogenic addiction (2), opioid addiction remains a major global public health and societal crisis (3). Addiction develops in a vulnerable subset of people (∼20%) who initiate opioid use recreationally and progressively transition to misuse (4–6). Despite decades of research, the determinants of vulnerability versus resilience to drug addiction remain largely unknown (6–8). The lack of a clear characterization of the differences between people who develop addiction and those who maintain controlled drug use continues to limit our understanding of the brain mechanisms of heroin addiction.

One key feature of addiction vulnerability is the progressive shift in reward valuation toward drugs and away from alternative reinforcers (9, 10). Consistent with this notion, a defining feature of addiction is a behavioral bias toward drug use, which drives people to prioritize the addictive drugs over healthier activities (11), including positive social interactions (12). Clinical studies suggest that this bias increases valuation of drug rewards, at the expense of natural rewards (9, 13–16). Drug valuation is accompanied by a shift toward drug-biased neural processing, characterized by increased dopamine release in striatal and limbic structures (17) and enhanced activation of prefrontal and limbic circuitry in response to drug-related stimuli compared to natural rewards (18–21). Importantly, neuroimaging studies directly comparing recreational users and people with addiction show that this exaggerated cue-induced prefrontal and limbic activation is specific to those with addiction and is not observed in recreational users (20, 22–24). These findings suggest that addiction involves neural adaptations that shift attention and motivation toward drug-related stimuli and away from natural rewards.

Preclinical models allow investigation of the neural mechanisms underlying addiction vulnerability (25). Studies show that ‘vulnerable’ rodents display distinct molecular and synaptic plasticity changes compared with ‘resilient’ rodents (21, 26–30). However, with few exceptions (31–33), researchers have identified vulnerable rodents solely based on drug-related behaviors and studied these behaviors in isolation using self-administration procedures where only the drug is available (34).

To address this gap, we recently developed a rat model of individual differences in heroin addiction (35) that incorporates a variation of a mutually exclusive choice between heroin and social interaction, previously shown to strongly inhibit opioid and psychostimulant self-administration (36–38). In this model, we first train rats to lever press for access to social interaction with a peer. We then trained them to lever press for heroin infusions without a post-infusion timeout. Next, we perform mutually exclusive choice sessions, allowing rats to choose between 5-min of social interaction vs. 5-min of heroin access (35). This procedure produces high variability in heroin intake, binge-like self-administration patterns, heroin seeking, and preference for heroin over social rewards, making it suitable to study brain mechanisms underlying opioid addiction vulnerability.

In the present study, we combined this behavioral procedure with positron emission tomography (PET) using 2-deoxy-2-[¹⁸F]fluoro-D-glucose (FDG), a marker of brain glucose metabolism (39, 40), to determine whether individual differences in ‘addiction severity’ are associated with distinct brain metabolic network states during prolonged abstinence at rest (homecage) and in response to cues previously associated with heroin or social self-administration. We used principal component analysis (PCA) to generate an addiction severity score based on heroin intake, binge-like episodes, and heroin preference.

## Materials and methods

### Subjects

We used 41 Sprague-Dawley rats (19 males and 22 females) and 41 social partner rats (Charles River; CD [SD], RRID:RGD_734476) weighing 125-150 g at arrival. We maintained the rats under reverse 12:12 h light/dark cycle (lights off at 8:00 A.M.) with food and water freely available. We housed two rats/cage prior to social self-administration and individually 3 days before social self-administration. We performed the experiment in accordance with the NIH Guide for the Care and Use of Laboratory Animals (8th edition), under a protocol approved by the local Animal Care and Use Committee. We excluded 5 of the 41 rats used in the study due to poor health (n=1), or issues during PET scan acquisition (n=4).

### Drugs

We received heroin hydrochloride (HCl) from the NIDA pharmacy and dissolved it in sterile saline. We chose a unit dose of 0.075 mg/kg for self-administration training based on our previous studies (35, 41).

### Intravenous surgery

We anesthetized the rats with isoflurane (5% induction, 2–3% maintenance, Covetrus). We attached silastic catheters to a modified 22-gauge cannula cemented to polypropylene mesh (Amazon or Industrial Netting), inserted the catheter into the jugular vein and fixed the mesh to the mid-scapular region of the rat (38, 42). We injected the rats with ketoprofen (2.5 mg/kg, s.c., Covetrus) after surgery and the following day to relieve pain and decrease inflammation. We allowed the rats to recover for 5-7 days before heroin self-administration training. During recovery and all experimental phases, we flushed the catheters every 24-48 h with gentamicin (4.25-5.0 mg/ml, Fresenius Kabi or Covetrus) dissolved in sterile saline. If we suspected catheter failure during training, we tested patency with Diprivan (propofol, NIDA pharmacy, 10 mg/mL, 0.1-0.2 mL injection volume, i.v.), and if not patent, we re-catheterized the other jugular vein and continued training after recovery.

### Self-administration chambers

Chambers were equipped with a stainless-steel grid floor, and two operant panels were placed on the left and right walls. The left panel was equipped with a houselight and the drug-paired active (retractable) lever. Responses on this lever activated the infusion pump and the discrete white-light cue located above the lever. The right panel was equipped with the social partner-paired active (retractable) lever. Responses on this lever activated the tone cue located above the lever and determined the opening of the guillotine-style sliding door.

#### Experimental procedures

The experiment included 7 phases: (1) social self-administration training, (2) heroin self-administration training plus 4 heroin-vs.-social interaction choice sessions, (3) early abstinence heroin-and social-seeking tests, (4) 7 consecutive days of heroin-vs.-social interaction choice sessions (5) homecage (at rest) PET imaging, and (6 and 7) late abstinence heroin- and social-seeking tests plus PET imaging (for experimental timeline see Figure 1).

**Figure 1.**
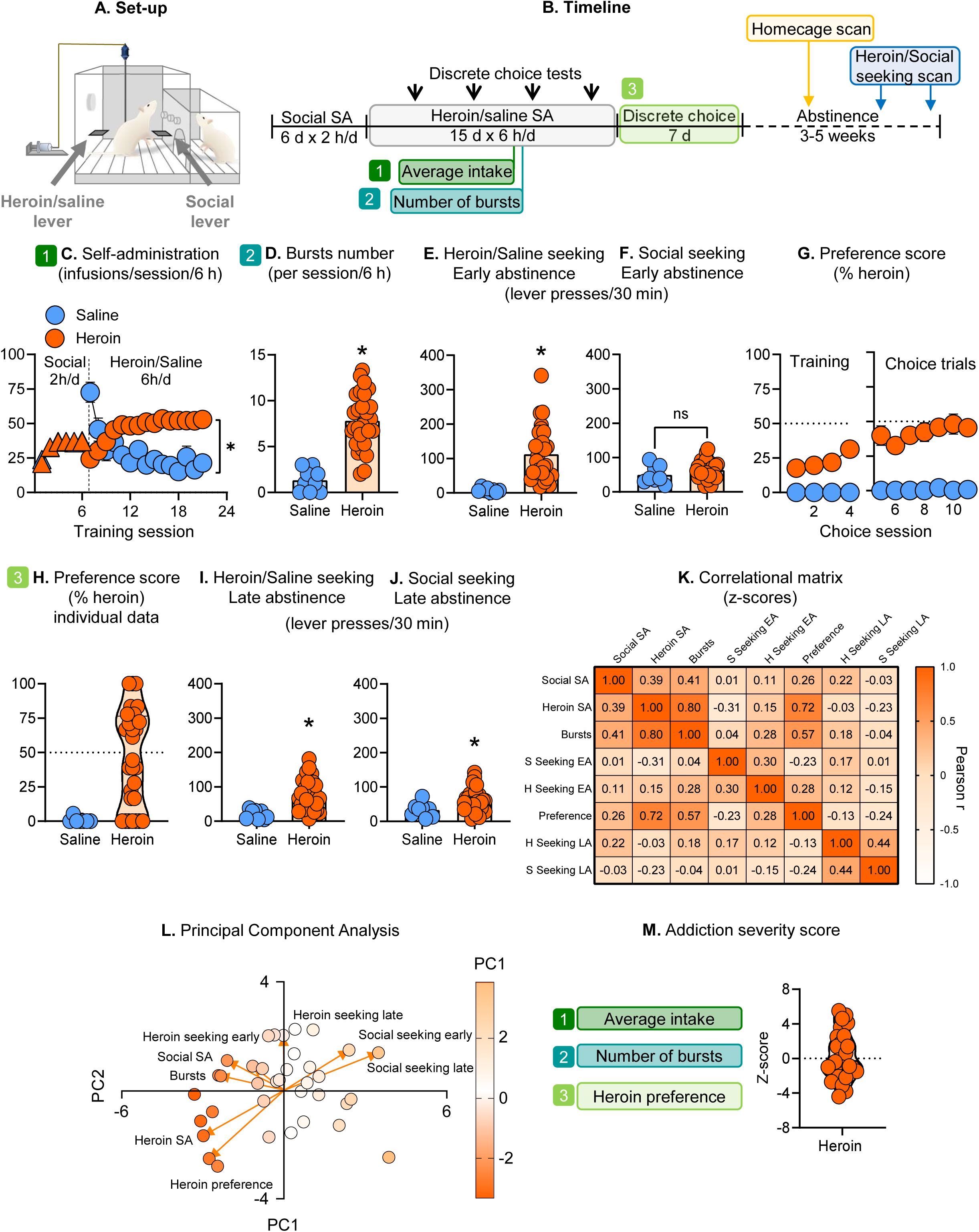
Multidimensional behavioral characterization of heroin addiction severity. (**A**) *Experimental setup*. (**B**) *Experimental timeline.* (**C**) *Social and heroin or saline self-administration*. Mean ± SEM number of social and heroin/saline rewards per session (saline: n = 8; heroin: n = 28). (**D**) *Bursts number*. Individual data of mean number of bursts during the last 3 sessions of heroin or saline self-administration training. (**E**) *Heroin- or saline-seeking tests – Early abstinence*. Individual data of number of lever presses on the active lever during the 30-minute extinction tests. (**F**) *Social-seeking test – Early abstinence.* Individual data of number of lever presses on the active lever during the 30-minute extinction tests. (**G***) Preference score during and after heroin self-administration training*. Mean ± SEM of percentage of heroin choices. 0 indicates a preference for social interaction and score of 100 indicates a preference for heroin. (**H**) *Preference score – Individual data*. Individual data of mean preference score during the last 3 sessions of choice. (**I**) *Heroin- or saline-seeking test – Late abstinence*. Individual data of number of lever presses on the active lever during the 30-minute extinction tests. (**J**) *Social seeking test – Late abstinence*. Individual data of number of lever presses on the active lever during the 30-minute extinction tests. (**K**) *Correlational matrix of behavioral measures of the heroin-trained rats.* (**I**) *Principal Component Analysis* (PCA) *of behavioral measures in heroin-trained rats.* (**M**) *Addiction severity score.* Individual data of severity z-score in heroin trained rats. * Different from saline, p<0.05.

### Social self-administration training

We trained rats to press a lever to have access to a social partner 2-h/d for 6 days. We housed the resident rats with their social partner (cage mate) until 3 days prior to social interaction self-administration; each resident rat lever pressed for its previously paired partner. The training sessions started with the illumination of the houselight and the insertion of the social partner-paired lever (that remained inserted throughout the session); responses on this lever resulted in the opening of the guillotine door and full-contact access to the social partner (1-min), paired with the tone cue (1-min) followed by a 2-min timeout (during which we manually placed the rats in their assigned compartment). In a pilot experiment, we observed that under both fixed ratio 1 (FR1) and progressive ratio reinforcement schedules, rats lever-pressed more for full-contact social interaction (see Supplementary Online Materials). Therefore, we allow the resident (self-administer) rats full physical access to their social partners during the training and choice sessions

### Social-seeking tests during early and later heroin abstinence

We tested the rats for social seeking under extinction conditions 1-3 days after heroin self-administration (early abstinence test) and 4-5 weeks later (late abstinence test). The test session (30 min) began with the extension of the active lever, and the illumination of the red houselight, which remained on for the duration of the session. Active lever presses during the social reward-seeking tests resulted in contingent presentations of the tone-cue and opening of the guillotine door, but no social partner was present in the social partner chamber.

### Drug self-administration training

After social self-administration, we trained the rats to self-administer heroin or saline 6-h/d for 15 days under continuous-access no-timeout conditions (41). Drug self-administration training sessions started with the insertion of the lever and the illumination of the houselight. Responses on the active lever (FR1 schedule) were reinforced by heroin infusion (3-s duration), paired with the cue light (3-s). After every three consecutive drug self-administration sessions, we tested rats for drug-vs.-social interaction preference in a discrete choice procedure (see below).

### Drug-seeking test early and late heroin abstinence

We tested the rats for social seeking under extinction conditions 1-3 days after heroin self-administration (early abstinence test) and 4-5 weeks later (late abstinence test). We counterbalanced the order of the heroin-and social seeking tests on abstinence days 1-2 and weeks 4-5.The duration of the test sessions was 30 min. The sessions began with the illumination of the houselight, followed 10-s later by the insertion of the drug-paired lever; the houselight remained on for the duration of the session. Lever presses during the tests resulted in the contingent presentation of the light cue (3-s), previously paired with heroin infusions, but no drug infusion was delivered. After the reward-seeking tests on days 1-3, we brought the rats to their homecages.

### Discrete choice

We assessed rat’s preference for heroin-vs.-social interaction in a discrete choice procedure. We conducted the choice procedure using the same parameters (dose of heroin and stimuli associated with the two active retractable levers) selected for the training phase. We allowed rats to choose between the heroin-paired and the social partner-paired levers. We tested rats for a total of 11 days. Choice sessions lasted for 6-h. Each 6-h choice session was divided into 6 discrete trials that were separated by 55-min. Each trial began with the flashing of the houselight for 30-s (a discriminative stimulus that signaled the choice session), followed by the illumination of the houselight and the insertion of both the social partner-paired and drug-paired levers. The rats then had to select one of the two levers. The operant response requirement for the lever’s selection was set to 5 consecutive responses (FR5) to avoid accidental choices. If the rats responded within 3-min, they had access to the selected lever for a total of 5-min under an FR1 reinforcement schedule. If the rats failed to respond on either active lever within 3-min, both levers were retracted, and the houselight was turned off with no reward delivery.

### Forced abstinence

We housed the rats in their homecages and handled them twice a week.

### Positron Emission Tomography (PET)

We assessed whole-brain metabolic activity, using FDG. FDG is a glucose analog that is taken up by metabolically-active cells and remains trapped for at least 60 min after uptake (43, 44), allowing to obtain a snapshot of regional metabolic activity during the awake, freely-moving state using PET.

We assessed whole-brain metabolic activity during abstinence at three different timepoints (Figure 1B): in the homecage (reference of resting brain metabolic activity), during heroin seeking, and during social seeking. We performed the scans one week apart and counterbalanced the rats for the heroin- and social-seeking tests. We focused our investigation during abstinence, to avoid potential confounds from the presence of heroin and its metabolites in the body or any additional metabolic changes induced by acute exposure to the drug. Additionally, in our previous study we showed that a key feature of addiction severity (i.e. the preference for heroin over social interaction) remained unchanged after 4 weeks of abstinence (41), indicating that the behavioral alterations persist.

Before the scans, we habituated the rats for three consecutive days prior to the scans to the transportation procedure, the PET facility environment, and the intraperitoneal (i.p.) injections with saline. Additionally, for the heroin- and social-seeking test scans, the day before the tests, we placed the rats in the “novel” self-administration chambers, for 10 minutes, immediately after an i.p. saline injection to acclimate them to the new environment. Rats were fasted for ∼18 h prior to PET imaging to minimize competition between circulating glucose and FDG and to ensure stable and reliable tracer uptake. <u>Note:</u> due to NIH safety rules, we could not test the rats in their original self-administration chambers.

On the days of the homecage scans, we injected each rat i.p. with ∼500-700 µCi (1 µCi/gram body weight) of FDG and then placed the rats in their homecage. After the uptake period, we anesthetized the rats with 2% isoflurane and positioned on a custom-made bed in a NanoScan small-animal PET/computed tomography (CT) scanner (Mediso Medical Imaging Systems). We scanned each rat for 30 min using a static imaging protocol (20-min), followed by a CT scan. <u>Note:</u> all rats were then fasted overnight (∼16 h) to attain consistency in blood glucose levels, as abnormal blood glucose levels interfere with FDG uptake (45).

On the days of the heroin- or social-seeking tests, we injected each rat i.p. with FDG and then we placed them in the self-administration chambers for a 30-min reward-seeking test (see above). After the uptake period, we anesthetized and scanned the rats.

We reconstructed PET data into static images using Mediso nanoScan© PET/CT software (Nucline, Version 3; Mediso, Budapest, Hungary). We applied a 3D iterative reconstruction algorithm (Tera-Tomo™ 3D), generating images with a 128 × 128 matrix and an isotropic voxel size of 0.4 mm. During reconstruction, we corrected for radioactive decay, PET dead time, random coincidences, and detector occupancy. We performed attenuation and scatter corrections using CT-based maps co-registered to the PET images, acquired immediately after the PET scan. We analyzed the images using the PMOD software environment (RRID:SCR_016547; PMOD Technologies).

### Statistical analysis

#### Behavioral data

We analyzed the data with the statistical programs SPSS (SPSS, Version 30), GraphPad Prism (Version 10) and XLSTAT. Before any analysis, we tested the data for sphericity and homogeneity of variance when appropriate. When the sphericity assumption was not met, we adjusted the degrees of freedom using the Greenhouse-Geisser correction. Outliers were included in the data analysis and presentation. The level of probability (p), for determining group differences, was set at p<0.05 (see Table S1, Supplementary Online Materials for complete reporting of the statistical analyses).

For the training phase, we analyzed the social self-administration data and the heroin/saline self-administration data using the within-subjects factor of Session and the between-subjects factor of Group (saline, heroin). For the choice tests, we calculated the preference (Preference Score) in the choice tests by calculating the percentage of heroin/saline choices relative to social choices (excluding omissions): number of heroin/saline choices / (heroin choices plus social choices) * 100. Then, we analyzed the preference score across sessions using the between-subjects factor of Group and the within-subjects factor of Session. For the heroin/saline seeking data, we analyzed the data using the between-subjects factor of Group and the within-subjects factor of Abstinence period (early, late). We also derived an ‘addiction severity score’ using principal component analysis (PCA). We calculated z-scores for the following variables: (1) mean social rewards, (2) mean heroin intake during the last 3 sessions of self-administration training, (3) number of bursts during the last 3 sessions of self-administration training [we define a burst as the self-administration of more than 3 infusions in less than 5 minutes (41)], (4) and (5) social and heroin seeking during early abstinence, (6) mean preference score during the last 3 sessions of choice training, (7) and (8) social and heroin seeking during late abstinence. Next, we calculated the ‘Addiction Severity’ score as the z-score sum of (2), (3) and (5).

#### PET data

We applied statistical parametric mapping analyses to the FDG-PET data in MATLAB R2023b using SPM12 (RRID:SCR_007037; https://www.fil.ion.ucl.ac.uk/spm/software/spm12/; University College London). Data were evaluated at p < 0.05 using the Probabilistic Threshold-Free Cluster Enhancement (pTFCE) method with multiple-comparisons correction (46) and a cluster extent threshold of 100 contiguous voxels (k = 100).

We first assessed differences in brain metabolic activity between saline-trained and heroin-trained rats. For this purpose, we performed voxel-based two-sample t-tests of whole-brain FDG uptake between heroin- and saline-trained rats for the homecage, heroin-seeking, and social-seeking scans. We then extracted voxel intensity from significant clusters and compared values between the two groups.

Next, we assessed whether FDG-PET imaging was sensitive to detecting differential metabolic activity patterns during the heroin- and social-seeking tests. For this purpose, we performed voxel-based paired t-tests of whole-brain FDG uptake between homecage scans and saline-, heroin-, or social-seeking scans in rats trained to self-administer either saline or heroin. We then extracted voxel intensity from significant clusters and compared it between homecage and test scans.

We next performed voxel-based regression analyses of whole-brain FDG uptake in heroin-trained rats for the homecage, heroin-seeking, and social-seeking scans using addiction severity scores as a regressor, to identify brain regions in which metabolic activity correlated with addiction severity. We extracted voxel intensity from significant clusters identified in the homecage scan, reflecting differences in FDG uptake at rest, and used these clusters as volumes of interest (VOIs) for correlational analyses.

## Results

### Behavioral data

#### Social and heroin self-administration

We first trained the rats for social and drug self-administration. During training, the rats increased the number of social rewards over time (main effect of Session, F _(2.41,82.1)_ = 33.96, p<0.001, Figure 1C). Next, we randomly assigned the rats to two groups: saline and heroin self-administration (matched for their social rewards). The rats increased the number of heroin, but not saline (Table S1), infusions earned over time (Session x Group interaction, F _(4.71,160.1)_ = 28.70, p<0.001, Figure 1C). The number of burst events was higher for heroin compared with saline (F _(1,34)_ = 30.43, p<0.001, Figure 1D).

#### Social- and heroin-seeking tests

Heroin seeking was higher in heroin-trained rats, compared with saline-trained rats, during both early and late abstinence (main effect of Group, F _(1,34)_ = 14.20, p<0.001, F _(1,34)_ = 4.78, p=0.036, Figure 1E, 1I). Social seeking did not significantly differ between saline- and heroin-trained rats during early abstinence (F _(1,34)_ = 132.73, p=0.170) but was higher in heroin-trained rats at late abstinence (main effect of Group, F _(1,34)_ = 4.78, p=0.036, Figure 1F, 1J).

#### Choice sessions

In the choice sessions, preference for heroin rewards was higher in heroin-trained compared with saline-trained rats (main effect of Group, F _(1,34)_ = 13.87, p<0.001*, Figure 1G), with significant individual variability in heroin preference (data not normally distributed) (Figure 1H).

#### Addiction severity score

PCA identified 3 main components (eigenvalue > 1; Figure 1L), which accounted for ∼71% of the total variance. The first principal component (eigenvalue 2.96; 34.2%), included variables related to self-administration and choice (Social Self-administration, Total Intake, Number of Bursts Events, and Heroin preference). The second component (eigenvalue 1.77; 20.5%), included variables related to late abstinence seeking (Late Abstinence Social- and Heroin-Seeking). The third component (eigenvalue 1.42; 16.4%), included variables related to early abstinence seeking (Early Abstinence Social- and Heroin-Seeking). Based on these results, we next derived a cumulative Severity z-score (Figure 1M) using heroin-specific behavioral measures only: heroin intake, number of burst events, and heroin preference - three variables that were highly intercorrelated (Figure 1K). Social self-administration was not included in the Severity z-score because it did not correlate with heroin preference in the choice test (Figure 1K), indicating that it reflects a behavioral aspect largely independent of heroin preference - a primary behavioral index of addiction-like behavior in our model. Additionally, removing social self-administration from the variable did not change the PCA results.

### PET data

#### Effects of heroin training on FDG uptake: heroin- vs. saline-trained rats (Figure 2)

Comparing homecage (at rest) FDG uptake during abstinence in saline- and heroin-trained rats we found that heroin abstinence significantly increases FDG uptake in a cluster of voxels (corrected p=0.05) including: motor cortex, insular cortex (p=0.008), somatosensory cortex (p=0.024), hippocampus (p=0.018), and pons (p=0.002) (Figure 2C-D). During heroin seeking, FDG uptake was significantly higher in heroin-compared to saline-trained rats in a cluster of voxels including the striatum (p=0.007), septum (p=0.008), thalamus (p=0.002), colliculus, and cerebellum (p<0.001) (Figure 2E-F). During social seeking, heroin self-administration, compared with saline, increased FDG uptake in a cluster of voxels including the retrosplenial cortex (p<0.001) and cerebellum (p<0.001) (Figure 2G-H).

**Figure 2.**
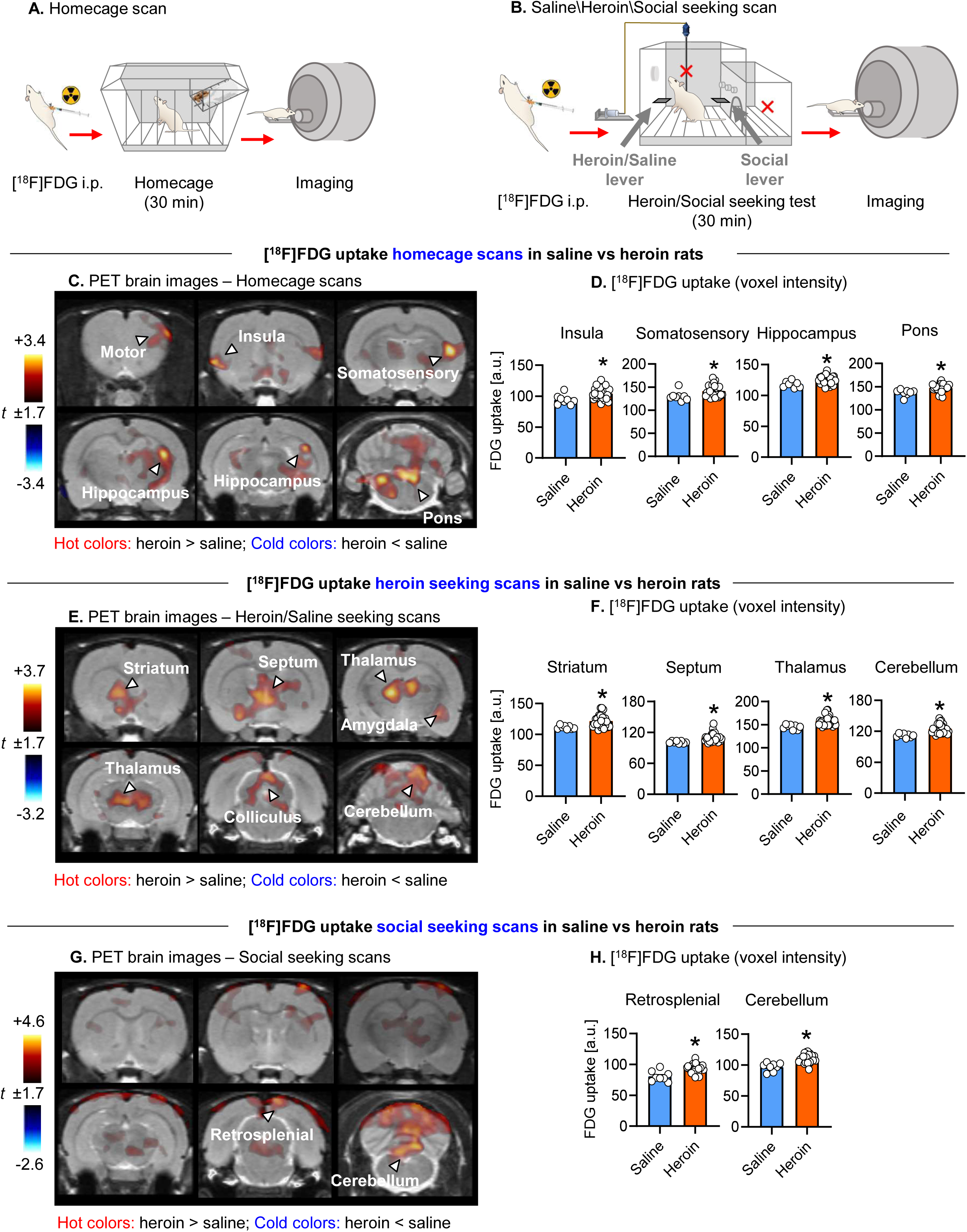
PET data: saline vs. heroin self-administration—homecage, heroin seeking, social seeking. (**A**) *Experimental timeline of homecage scans*. (**B**) *Experimental timeline of reward-seeking scans*. (**C**) *PET brain images homecage scan*. Coronal PET brain images overlaid on structural MRI showing significant differences in [^18^F]FDG uptake in homecage scans in control, saline-trained rats (n = 8 rats) and heroin-trained rats (n = 28 rats) (two-sample t-test, saline vs heroin) [Hot colors: heroin > saline; cold colors: heroin < saline] (cluster corrected, p < 0.05, minimum cluster size = 100 voxels). (**D**) *[^18^F]FDG uptake in insula, somatosensory cortex, hippocampus, and pons uptake –* Individual normalized FDG uptake values (arbitrary units [a.u.] of voxel intensity) extracted from significant clusters (defining *insula, S1, hippocampus or pons*) showing significant differences between control saline- and heroin-trained rats in homecage scans. (**E**) *PET brain images heroin- or saline-seeking scans*. Coronal PET brain images overlaid on structural MRI showing significant differences in [^18^F]FDG uptake in heroin/saline seeking scans in control, saline-trained rats (n = 8 rats) and heroin-trained rats (n = 28 rats) (two-sample t-test, saline vs heroin). [Hot colors: heroin > saline; cold colors: heroin < saline] (cluster corrected, p < 0.05, minimum cluster size = 100 voxels). (**F**) *[^18^F]FDG uptake in striatum, septum, thalamus, and cerebellum uptake –* Individual normalized FDG uptake values (arbitrary units [a.u.] of voxel intensity) extracted from significant clusters (defining striatum, septum, thalamus, and cerebellum) showing significant differences between control saline- and heroin-trained rats in homecage scans. (**G**) *PET brain images social-seeking scans*. Coronal PET brain images overlaid on structural MRI showing significant differences in [^18^F]FDG uptake in social seeking scans in control, saline-trained rats (n = 8 rats) and heroin-trained rats (n = 28 rats) (two-sample t-test, saline vs heroin). [Hot colors: heroin > saline; cold colors: heroin < saline] (cluster corrected, p < 0.05, minimum cluster size = 100 voxels). (**H**) *[^18^F]FDG uptake in retrosplenial and cerebellum –* Individual normalized FDG uptake values (arbitrary units [a.u.] of voxel intensity) extracted from significant clusters (defining retrosplenial or cerebellum) showing significant differences between control saline- and heroin-trained rats in social seeking scans. Abbreviations: a.u. = arbitrary units.

#### FDG uptake during heroin and social seeking in heroin-trained rats: comparison with homecage uptake (Figure 3)

We compared FDG uptake during homecage scans with FDG uptake during heroin and social seeking scans in heroin-trained rats. Overall brain metabolic activity patterns differed between heroin and social seeking when comparing homecage and the reward-seeking test scans (Figure 3C and 3E). Both heroin and social seeking, compared to homecage conditions, were associated with reduced FDG uptake in the in the olfactory bulb (OB) (heroin seeking p<0.001; social seeking p<0.001) (Figure 3C-F). Heroin seeking significantly increased FDG uptake in clusters of voxels in different brain regions including the olfactory tubercule (OT), motor cortex, hypothalamus (p= p<0.001), ventral hippocampus (vHipp), reticular formation, and cerebellum (p<0.001) (Figure 3C-D). Social seeking significantly increased FDG uptake in clusters of voxels in different brain regions including the OT, motor cortex, different areas of the thalamus, retrosplenial cortex, and cerebellum (p<0.001).

**Figure 3.**
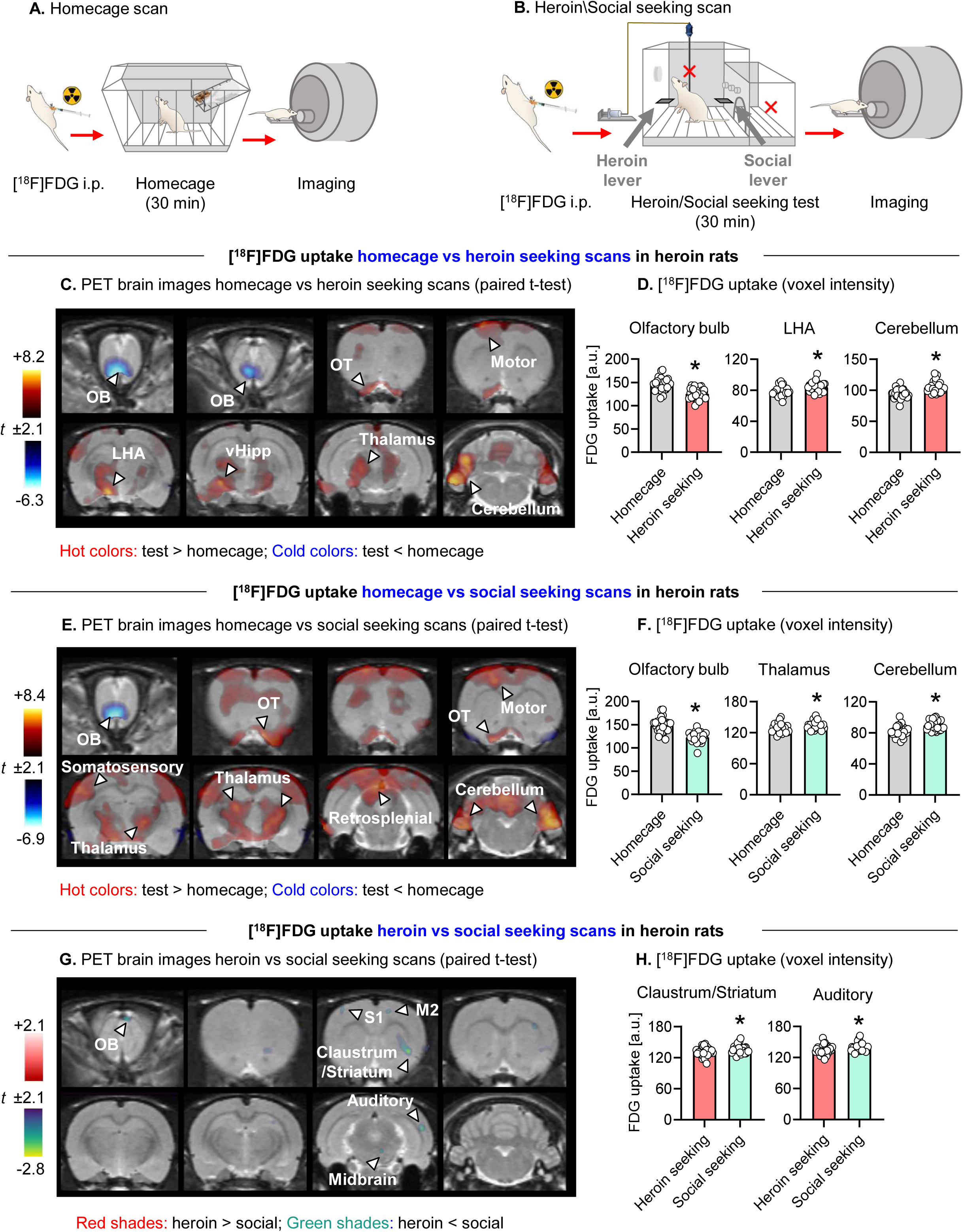
PET data: homecage vs. heroin seeking or social seeking in heroin-trained rats. (**A**) *Experimental timeline of homecage scans*. (**B**) *Experimental timeline of seeking scans*. (**C**) *PET brain images homecage vs heroin-seeking scan*. Coronal PET brain images overlaid on structural MRI showing significant differences in [^18^F]FDG uptake in homecage vs heroin seeking scans in heroin-trained rats (n = 28 rats) (paired t-test, homecage vs heroin seeking). [Hot colors: test > homecage; cold colors: test < homecage] (cluster corrected, p < 0.05, minimum cluster size = 100 voxels). (**D**) *[^18^F]FDG uptake in olfactory bulb, hypothalamus, and cerebellum –* Individual normalized FDG uptake values (arbitrary units [a.u.] of voxel intensity) extracted from significant clusters (defining olfactory bulb, hypothalamus, or cerebellum) showing significant differences between homecage scan and heroin-seeking scan in heroin-trained rats. (**E**) *PET brain images homecage/social seeking scan*. Coronal PET brain images overlaid on structural MRI showing significant differences in [^18^F]FDG uptake in homecage vs social seeking scans in heroin-trained rats (n = 28 rats) (paired t-test, homecage vs social seeking). [Hot colors: test > homecage; cold colors: test < homecage.] (cluster corrected, p < 0.05, minimum cluster size = 100 voxels). (**F**) *[^18^F]FDG uptake in olfactory bulb, thalamus, and cerebellum –* Individual normalized FDG uptake values (arbitrary units [a.u.] of voxel intensity) extracted from significant clusters (defining olfactory bulb, thalamus, or cerebellum) showing significant differences between homecage scan and social seeking scan in heroin-trained rats (n = 28 rats). (**G**) *PET brain images heroin- and social-seeking scans*. Coronal PET brain images overlaid on structural MRI showing significant differences in [^18^F]FDG uptake in heroin vs social seeking scans in heroin-trained rats (n = 28 rats) (paired t-test, heroin seeking vs social seeking). [Red shades: heroin > social; Green shades: heroin < social] (cluster corrected, p < 0.05, minimum cluster size = 100 voxels) (paired t-test, heroin seeking vs social seeking). (**H**) *[^18^F]FDG uptake in claustrum and auditory cortex –* Individual normalized FDG uptake values (arbitrary units [a.u.] of voxel intensity) extracted from significant clusters (defining cluastrum or auditory cortex) showing significant differences between heroin seeking scans and social seeking scans in heroin-trained rats. Abbreviations: a.u. = arbitrary units; OT = olfactory tubercule; OB = olfactory bulb; M2 = secondary motor cortex; S1 = primary somatosensory cortex.

We also directly compared heroin- and social-seeking scans. We found that compared to social seeking, heroin seeking was associated with decreased FDG uptake in a cluster of voxels including motor cortex, somatosensory cortex, claustrum (p<0.001), midbrain and auditory cortex (p=0.003) (Figure 3G-H).

#### Association between FDG uptake and addiction severity in heroin-trained rats (Figure 4)

To test whether addiction severity is related to alterations in brain metabolism, we performed a regression analysis of whole brain FDG uptake using the addiction severity score as a regressor. At rest scan, higher addiction severity was associated with reduced FDG uptake in the piriform cortex (PirCx) (Pearson *r =* -0.39, p=0.038) and increased FDG uptake in the vHipp (Pearson *r =* 0.55, p=0.002, Figure 4B-D). At the heroin-seeking scan, higher addiction severity was associated with reduced FDG uptake in clusters of voxels in the postsubiculum (Pearson *r = -*0.65, p<0.001) and cerebellum (Pearson *r = -*0.53, p=0.003) and lower effect in the striatum (Pearson r = -0.40, p=0.03) (Figure 4E-G). At the social-seeking scan, higher addiction severity was associated with reduced FDG uptake in the visual cortex (Pearson -*r =* 0.49, p=0.008) (Figure 4H-I).

**Figure 4.**
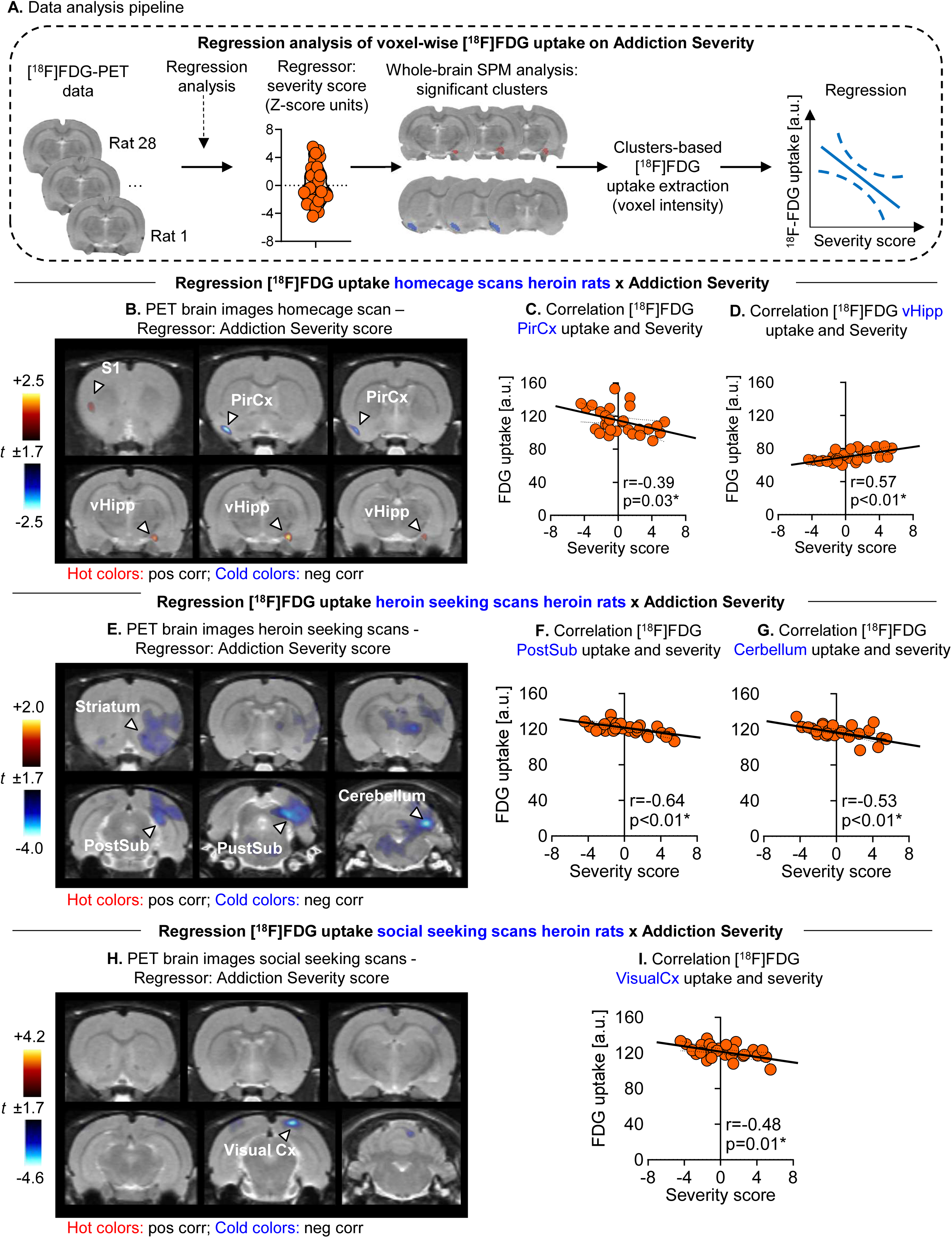
Correlations between addiction severity and FDG uptake at homecage and during heroin or social seeking. (**A**) *Data analysis pipeline*. Sequential steps for regression analysis of FDG uptake on Addiction Severity. (**B**) *PET brain images homecage scan – Regressor: Addiction Severity score.* Coronal PET brain images overlaid on structural MRI showing cluster of voxels in which FDG uptake at rest was significantly associated with addiction severity in heroin-trained rats (n = 28 rats). [Hot colors: positive association between FDG uptake and severity; cold colors: negative association between FDG uptake and severity.] (cluster corrected, p < 0.05, minimum cluster size = 100 voxels). (**C**) *Correlation FDG PirCx uptake and Addiction Severity. (**D**) Correlation [^18^F]FDG vHipp uptake and Addiction Severity.* (**E**) *PET brain images brain images heroin-seeking scans – Regressor: Addiction Severity score.* Coronal PET brain images overlaid on structural MRI showing cluster of voxels in which FDG uptake during heroin seeking was significantly associated with addiction severity in heroin-trained rats (n = 28 rats). [Hot colors: positive association between FDG uptake and severity; cold colors: negative association between FDG uptake and severity.] (cluster corrected, p < 0.05, minimum cluster size = 100 voxels). (**F**) *Correlation [^18^F]FDG PostSub uptake and Addiction Severity. (**G**) Correlation FDG Cerebellum uptake and Addiction Severity.* (**H**) *PET brain images brain images social seeking scans – Regressor: Addiction Severity score.* Coronal PET brain images overlaid on structural MRI showing cluster of voxels in which FDG uptake during social seeking was significantly associated with addiction severity in heroin-trained rats (n = 28 rats). [Hot colors: positive association between FDG uptake and severity; cold colors: negative association between FDG uptake and severity.] (cluster corrected, p < 0.05, minimum cluster size = 100 voxels). (**I**) *Correlation FDG Visual cortex (VisualCx) uptake and Addiction Severity.* Abbreviations: a.u. = arbitrary units; PirCx = piriform cortex; PostSub = Post subiculum S1 = primary somatosensory cortex; vHipp = ventral hippocampus; VisualCx = visual cortex.

#### FDG uptake in low vs. high addiction severity

To further investigate the relationship between brain metabolism and addiction severity, we performed a two-sample t-test of whole-brain FDG uptake in rats classified as highly addicted (top quartile) compared with low-addicted rats (bottom quartile) (Figure 5A). The two-sample t-test performed on the homecage scans confirmed reduced FDG uptake in the PirCx (p=0.001) and increased FDG uptake in the vHipp (p<0.001). Additionally, this analysis showed increased metabolic activity in the nucleus accumbens (NAc, p=0.017) and lateral hypothalamus (LHA, (p=<0.001)). The same analysis during heroin seeking showed reduced FDG uptake in the postsubiculum (p=0.03), and cerebellum (p=0.002). Notably, we didn’t observe any effect during social seeking.

**Figure 5.**
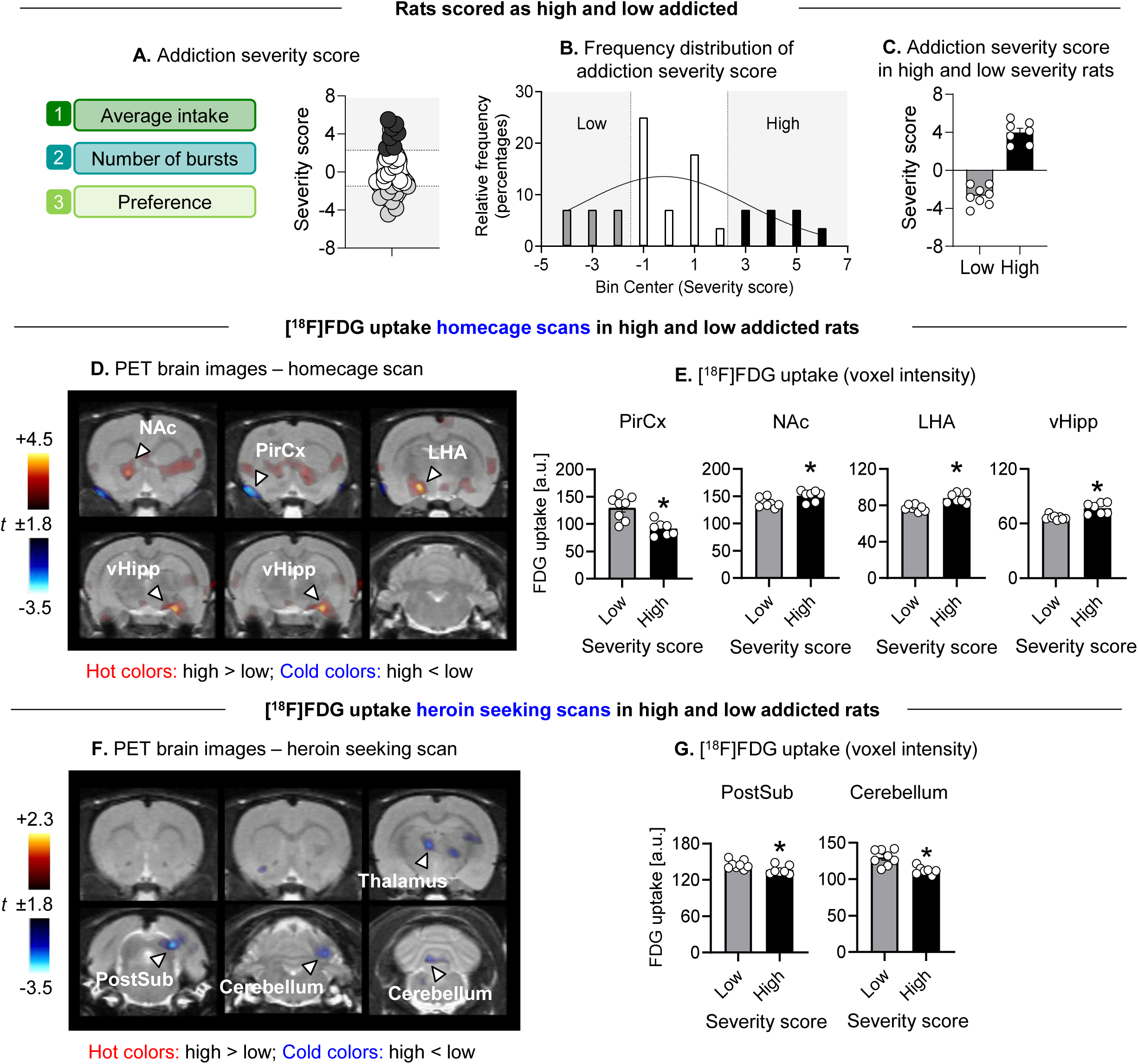
FDG uptake at homecage and during heroin seeking in rats with low and high addiction score. (**A**) *Addiction severity score*. (**B**) *Frequency distribution of addiction severity score* (**C**) *Addiction severity score in high (upper quartile) and low (lower quartile) addiction severity rats.* (**D**) *PET brain images - homecage scan*. Coronal PET brain images overlaid on structural MRI showing significant differences in [^18^F]FDG uptake in homecage scans in rats classified as low (n = 8) and high (n = 7) addiction severity. [Hot colors: high > low; cold colors: high < low.] (cluster corrected, p < 0.05, minimum cluster size = 100 voxels). (**E**) *[^18^F]FDG uptake in PirCx, LHA, and vHipp*. Individual normalized FDG uptake values (arbitrary units [a.u.] of voxel intensity) extracted from significant clusters (defining PirCx, NAc, LHA or vHipp) showing significant differences between low- and high-addiction–scored rats in homecage scans. (**F**) *PET brain images - homecage scan*. Coronal PET brain images overlaid on structural MRI showing significant differences in [^18^F]FDG uptake in heroin seeking scans in low addiction scored (n = 8 rats) and high addiction scored rats (n = 7 rats). [Hot colors: high > low; cold colors: high < low.] (cluster corrected, p < 0.05, minimum cluster size = 100 voxels). (**G**) *[^18^F]FDG uptake in PostSub and Cerebellum*. Individual normalized FDG uptake values (arbitrary units [a.u.] of voxel intensity) extracted from significant clusters (defining *PostSub or Cerebellum*) showing significant differences between low- and high-addiction–scored rats in homecage scans. Abbreviations: a.u. = arbitrary units; NAc = nucleus accumbens, LHA = lateral hypothalamus; PirCx = piriform cortex; PostSub = post subiculum; vHipp = ventral hippocampus. *Different from low, p<0.05.

## Discussion

We identified distinct metabolic alterations associated with heroin self-administration, prolonged abstinence, and addiction vulnerability. Three principal observations emerged. First, prolonged abstinence following heroin self-administration was associated with widespread increases in FDG uptake across cortical and subcortical regions, suggesting a persistent shift in basal brain metabolic activity. Second, heroin-trained rats showed elevated heroin seeking during both early and late abstinence. In contrast, despite the metabolic alterations at rest, social seeking remained intact: it was comparable to controls during early abstinence and increased during late abstinence. Addiction severity was not correlated with social seeking at either time point. Moreover, although heroin and social seeking engaged overlapping brain regions, social seeking produced comparable metabolic activation in both saline- and heroin-trained rats. Together, these observations suggest drug-specific adaptations rather than deficits in natural rewards processing. Third, addiction severity was associated with distinct metabolic alterations that were most pronounced under resting conditions. Higher severity was associated with reduced piriform cortex and increased ventral hippocampal metabolism at rest, whereas during heroin seeking, addiction severity was associated with reduced activity of postsubiculum and cerebellum.

### Heroin self-administration and vulnerability to heroin addiction are associated with drug-specific brain metabolic alterations without natural reward deficits

In prior work, we showed that heroin preference over social interaction remains stable after four weeks of abstinence, suggesting a persistent shift in the relative value of heroin over social interaction (35). Based on this and prior clinical (18, 47–50) and preclinical (51–55) evidence suggesting enhanced drug salience and blunted responses to natural rewards in addiction, we hypothesized that prolonged abstinence would be associated with heightened neural responses to heroin cues and reduced responses to social cues during the relapse tests.

Our findings did not support this hypothesis. Behaviorally, social seeking was comparable between the heroin- and saline-trained rats during early abstinence and was even elevated in heroin-trained rats during late abstinence. Moreover, addiction severity did not correlate with social-seeking behavior. Neuroimaging similarly showed minimal differences between groups during social seeking, and high- and low-severity rats did not differ in FDG uptake during social seeking. Together, these results indicate that neither heroin self-administration nor addiction severity was associated with deficits in social behavior in our model.

These results are consistent with previous findings from our lab (56) and others (57, 58) showing intact sociability and operant responding for social rewards during protracted withdrawal from non-contingent opioid exposure in rats. However, they contrast with previous preclinical and clinical evidence, as well as addiction theories, suggesting that opioid exposure results in generalized anhedonia, reduced sensitivity to natural rewards (54, 59) and impairments in social functioning (53, 60, 61). Although such deficits are frequently observed in individuals with opioid addiction, it remains unclear whether they reflect preexisting vulnerability, consequences of prolonged drug exposure, or comorbid psychiatric conditions (62, 63). Indeed, recent clinical studies report heightened behavioral and neural responses to drug cues in individuals with addiction without consistent evidence for global reward deficits, or generalized anhedonia, suggesting that the capacity to experience pleasure from natural rewards is preserved (9, 18, 64, 65).

Although the present findings agree with recent clinical evidence, important questions remain. Future studies are needed to further characterize potential anhedonic phenotypes in our model, particularly during early abstinence. Indeed, acute opioid withdrawal is associated with transient suppression of response to natural rewards (56, 66), and these early adaptations were not investigated in our study.

### Neurometabolic signature of heroin addiction vulnerability

The primary objective of this study was to characterize the neurometabolic signature of addiction vulnerability by examining whether individual differences in heroin addiction severity are associated with differential metabolic activity during heroin and social seeking.

Contrary to our expectations, we found that higher addiction severity was primarily associated with differences in metabolic activity at rest, including increased ventral hippocampus metabolism and reduced piriform cortex metabolism. During heroin seeking, higher addiction severity was associated with reduced activity of postsubiculum and cerebellum. These findings suggest that addiction vulnerability in our model is characterized predominantly by metabolic alterations at rest rather than alterations in reward seeking-dependent activation, potentially reflecting pre-existing behavioral and genetic vulnerability to drug self-administration (67–74). Future studies should investigate the temporal dynamics of these adaptations to determine whether they represent a stable trait that predispose individuals to heroin-taking or reflect persistent neurobiological changes that emerge after drug self-administration [e.g., (26, 75)].

The shifts in metabolic activity at rest are consistent with prior evidence of persistent cellular and synaptic adaptations in addiction-vulnerable rodents (26, 28, 29, 31, 76). However, while these findings suggest that stable resting-state differences bias neural responses to drug-associated cues, our imaging approach does not allow direct assessment of circuit-level recruitment. Future studies using circuit-specific, causal, and high-resolution methods are necessary to determine whether baseline alterations in each node (e.g., ventral hippocampus or piriform cortex) differentially recruit downstream brain regions in high- versus low- addiction severity rats.

Finally, the regions showing basal alterations in our study are consistent with prior evidence implicating the ventral hippocampus and its projections to the prefrontal cortex and nucleus accumbens in reward seeking and addiction- and relapse-related behaviors across drug classes (77–82). Similarly, the piriform cortex, and its connections to orbitofrontal cortex, anterior insular cortex, and amygdala, have been linked to relapse to opioid seeking (83–85) and opioid-induced social behavior changes (86).

### Abstinence-related metabolic alterations following heroin self-administration

Heroin abstinence was associated with widespread increases in brain FDG uptake. This hypermetabolic pattern contrasts with prior clinical and preclinical FDG-PET studies reporting hypometabolism during opioid abstinence (87–89). However, methodological factors likely contribute to these differences.

In rats, Soto-Montenegro et al. (87) reported widespread reduction in FDG uptake 14 days after morphine self-administration. However, the authors used a different behavioral procedure and morphine instead of heroin. Given the evidence that distinct self-administration schedules produce divergent neurobiological adaptations (90, 91) and the pharmacological differences between morphine and heroin (92), these methodological factors may underlie the discrepant findings. Furthermore, Soto-Montenegro et al. (87) observed strain-dependent metabolic effects (including between saline controls). Considering that they used Lewis and F344 rats, whereas we used Sprague–Dawley rats, and the evidence that genetic background influences addiction vulnerability and drug-induced neuroplasticity (30, 93, 94), strain differences likely contribute to the divergent findings.

Clinical studies similarly report hypometabolism in opioid-dependent individuals (88, 89); however, participants are typically maintained on methadone, previously treated with methadone, or are polysubstance users, precluding direct comparison with our findings.

### Methodological considerations

We used FDG PET to assess brain glucose metabolism as a proxy for brain activity. Although this approach represents a major strength, such as allowing whole-brain and behavior-related metabolic activity (44), it also has interpretational limitations.

In several instances, we observed reductions in FDG uptake that were counterintuitive based on prior literature using techniques that more directly assess neuronal activation. For example, previous studies investigating individual differences in addiction vulnerability, particularly those using immediate early genes such as Fos (95), have often reported increased neuronal activation in specific brain regions during addiction-related behaviors (28, 96). Thus, we expected increased brain activity in heroin-vulnerable rats during heroin seeking; instead, we found reduction in FDG uptake in the postsubiculum, suggesting decreased metabolic activity.

Accumulating evidence indicates that FDG uptake does not exclusively reflect neuronal activity (97), and the cellular sources of the FDG signal and the mechanisms regulating its variations are still debated. Indeed, studies show that FDG is preferentially taken up by glial cells: Compared with neurons, uptake is about twofold higher in astrocytes and threefold higher in microglia (98). These findings suggest that changes in FDG signal cannot be interpreted solely as changes in neuronal activity. Rather, it reflects integrated metabolic activity across neurons and glial cells, including astrocytes and microglia. Thus, reduced FDG uptake in vulnerable rats does not necessarily indicate decreased neuronal activity and impaired functioning, but may instead reflect alterations in glial metabolism, neuroimmune signaling, or neuron-glia communication (97, 98). Future studies combining FDG imaging with established measures of neuronal activation, such as Fos mapping, cell-type specific calcium imaging or electrophysiology, are needed to determine how metabolic changes relate to neuronal activity in addiction vulnerability.

## Conclusions and clinical implications

We found that heroin seeking and addiction severity were associated with distinct metabolic patterns across multiple brain regions, including areas not traditionally linked to opioid addiction such as the postsubiculum and cerebellum. These findings identify neurometabolic features of heroin addiction vulnerability in a rat model that may provide biomarkers for identifying opioid addiction risk, monitoring treatment responses, and informing relapse-prevention therapies. In contrast, heroin self-administration and addiction severity did not alter social-seeking behavior or its associated brain metabolic activity, suggesting that sensitivity to social reward remains preserved in our opioid addiction model. This observation is consistent with recent preclinical (56–58) and clinical studies (9, 18, 64, 65). Preserved responses to social rewards support social-based behavioral treatments, such as the community reinforcement approach, which leverages social reinforcement to promote abstinence (99).

---

**Figure S1.**
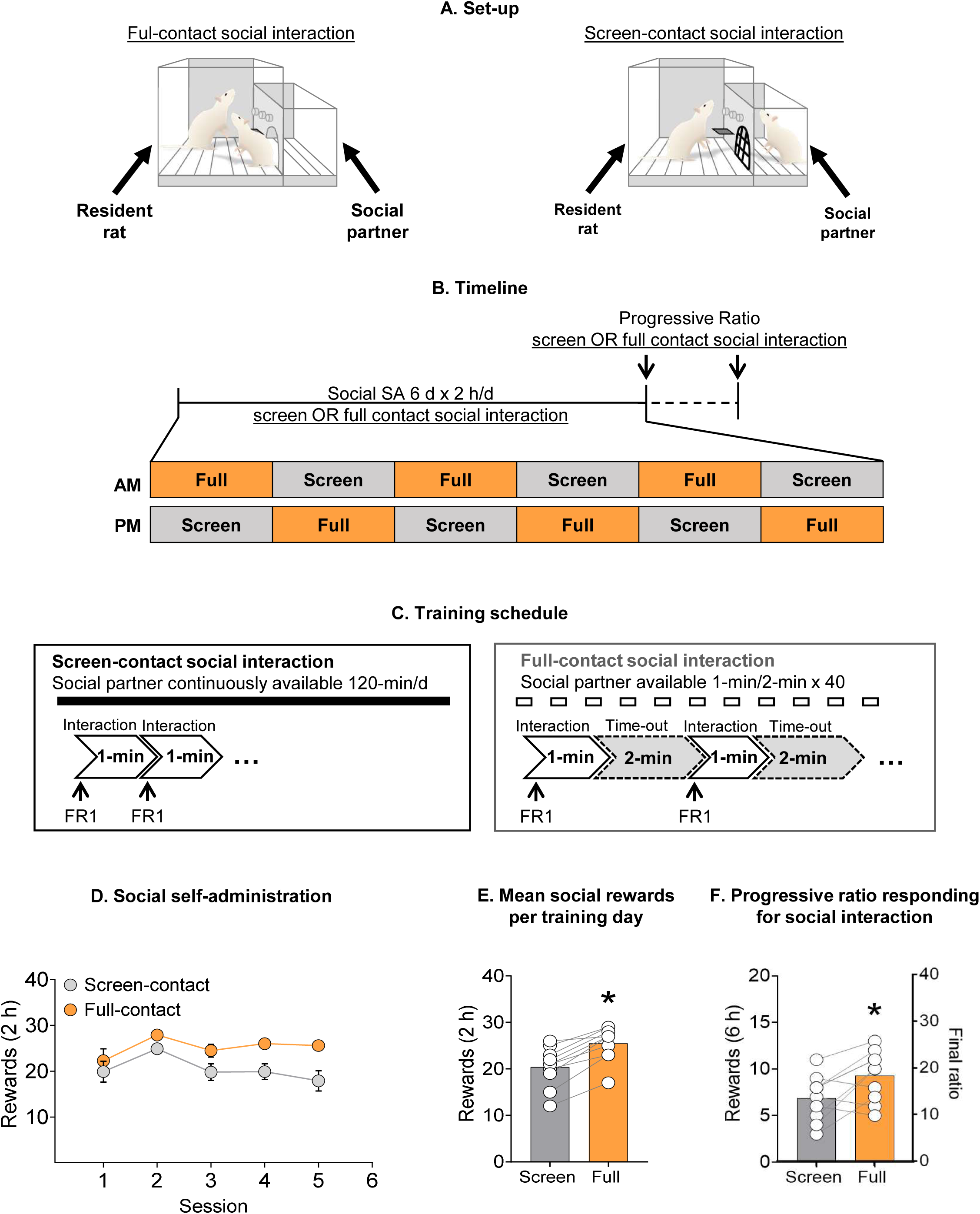
Social self-administration: full social contact versus limited (screen) social contact. **(A)** *Experimental setup*. **(B)** *Experimental timeline.* **(C)** *Experimental training schedule.* **(D)** *Social self-administration*. Mean ± SEM number of full contact or screen-contact social rewards per session. (n = 10, within-subjects). **(E)** *Mean rewards for social self-administration*. Individual data of mean number of social rewards though all sessions of full/screen-contact social self-administration training. **(F)** *Progressive ratio responding*. Individual data of social rewards earned during a single progressive ratio session for full-contact and screen-contact social self-administration. Left y-axis indicates rewards obtained and right y-axis indicates final-ratio requirement achieved.

**Figure S2.**
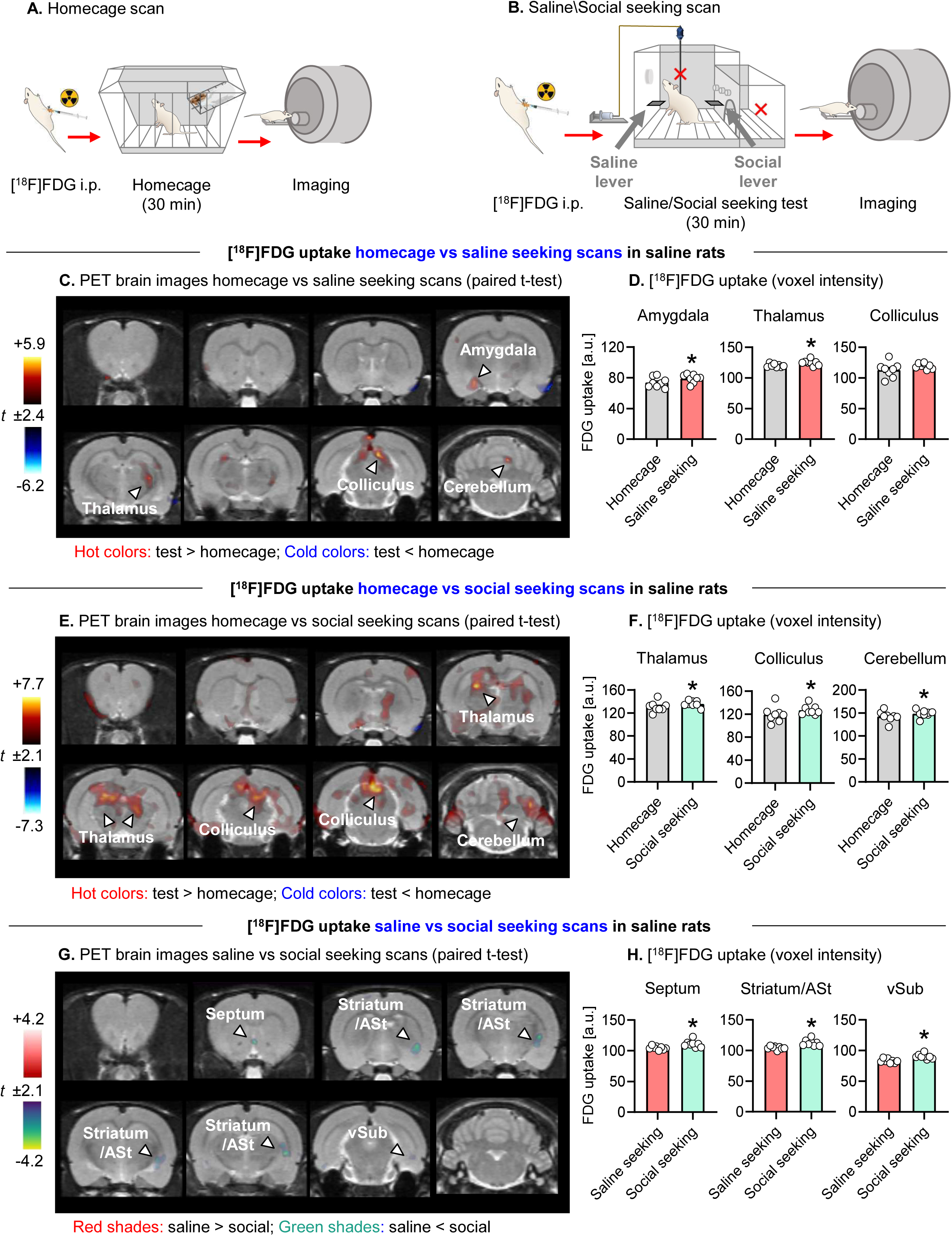
PET data: homecage vs. heroin seeking or social seeking in saline-trained rats. **(A)** *Experimental timeline of homecage scans*. **(B)** *Experimental timeline of seeking scans*. **(C)** *PET brain images homecage vs saline-seeking scan*. Coronal PET brain images overlaid on structural MRI showing significant differences in FDG uptake in homecage vs saline-seeking scans in saline-trained rats (n = 8 rats) (paired t-test, homecage vs saline seeking). [Hot colors: test > homecage; cold colors: test < homecage] (cluster corrected, p < 0.05, minimum cluster size = 100 voxels). **(D)** *FDG uptake in amygdala, thalamus, and colliculus –* Individual normalized FDG uptake values (arbitrary units [a.u.] of voxel intensity) extracted from significant clusters (defining amygdala, thalamus, or colliculus) showing significant differences between homecage scan and saline-seeking scan in saline-trained rats. **(E)** *PET brain images homecage/social seeking scan*. Coronal PET brain images overlaid on structural MRI showing significant differences in FDG uptake in homecage vs social-seeking scans in saline-trained rats (n = 8 rats) (paired t-test, homecage vs social seeking). [Hot colors: test > homecage; cold colors: test < homecage.] (cluster corrected, p < 0.05, minimum cluster size = 100 voxels). **(F)** *FDG uptake in thalamus, colliculus, and cerebellum –* Individual normalized FDG uptake values (arbitrary units [a.u.] of voxel intensity) extracted from significant clusters (defining thalamus, colliculus, or cerebellum) showing significant differences between homecage scan and social seeking scan in saline-trained rats.**(G)** *PET brain images saline/social seeking scan*. Coronal PET brain images overlaid on structural MRI showing significant differences in FDG uptake in saline vs social-seeking scans in saline-trained rats (n = 8 rats) (paired t-test, saline seeking vs social seeking). [Red shades: heroin > social; Green shades: heroin < social] (cluster corrected, p < 0.05, minimum cluster size = 100 voxels) (paired t-test, saline seeking vs social seeking). **(H)** *FDG uptake in septum, striatum, and vSub –* Individual normalized FDG uptake values (arbitrary units [a.u.] of voxel intensity) extracted from significant clusters (defining septum, striatum, or vSub) showing significant differences between saline seeking scans and social seeking scans in saline-trained rats. Abbreviations: a.u. = arbitrary units; vSUb = Ventral subiculum.

## Supplementary Online Materials

### Social self-administration: full social contact versus limited (screen) social contact Materials and methods

#### Subjects

We used a total of 10 Sprague–Dawley rats (4 males and 6 females). Males weighed 400–500 g and females weighed 300–350 g.

#### Self-administration chambers

We equipped the chambers with a stainless-steel grid floor and installed two operant panels on the left and right walls. The left panel contained a houselight and a drug-paired active (retractable) lever.

Responses on this lever, which we did not use in the current study, activate the infusion pump and the discrete white-light cue located above the lever. The right panel contained the social partner-paired active (retractable) lever. Responses on this lever activated a tone cue located above the lever and triggered the opening of a guillotine-style sliding door. We assigned each rat to two distinct operant chambers: one configured for partial (screen) social-contact access, equipped with a stainless-steel grid screen that allowed visual, olfactory, and limited tactile interaction after the opening of the guillotine door, and another configured for full-contact access, in which the opening of the guillotine door allowed free movement between compartments for unrestricted social interaction.

#### Social self-administration

We trained rats to press a lever for access to a social partner for 2 h/day, divided into two daily sessions (morning and afternoon), over 5 days. Each rat completed one full-contact and one screen-contact session per day in two different cages, with session order counterbalanced across subjects. Resident rats (lever pressers) were single-housed, and we assigned each rat a novel conspecific as a social partner. These social partners had previously participated as partners in earlier experiments. Each resident rat remained paired with the same social partner throughout the experiment. Each training session began with illumination of the house light and insertion of the social partner–paired lever, which remained extended for the duration of the session. During full-contact access sessions, responses on the active lever were reinforced under a fixed ratio 1 (FR-1) reinforcement schedule and resulted in the opening of the guillotine door and full-contact access to the social partner (1-min), paired with the tone cue (1-min) followed by a 2-min timeout (during which the rats were manually placed back to their assigned compartment) (Figure S1C). During screen-contact access sessions, responses on the active lever were reinforced under anFR-1 reinforcement schedule and resulted in the opening of the guillotine door and screen-contact with the social partner social partner (1-min), paired with the tone cue (1-min) followed by no timeout (Figure S1C). In both conditions, we set a maximal number of rewards to 40.

#### Social self-administration under a progressive ratio schedule

After completing social self-administration training under an FR1 reinforcement schedule, we tested rats for social self-administration under a progressive ratio (PR) reinforcement schedule (1). In this schedule, the response requirement increased according to the formula (5 × e^(0.2 × n)) − 5, where *n* represents the position of the ratio in the sequence. This progression results in the following response requirements for successive reinforcers: 1, 2, 4, 6, 9, 12, 15, 20, 25, 32, 40, etc. We conducted two separate PR test sessions to assess responding for full- and screen-contact social access (1-min), with the order of testing counterbalanced across subjects so that each rat experienced both conditions. Each 6-h test session began with extension of the active lever and illumination of the house light, which remained on for the duration of the session. Active lever presses resulted in contingent presentation of the tone cue, opening of the automatic guillotine door for 1-min, and full- or screen-contact social access to the social partner.

### Statistical analysis

For the training phase, we analyzed the social self-administration data using the between-subjects factor of Access Condition (screen-contact, full-contact) and the within-subjects factor of Session. For the progressive-ratio test, we analyzed the number of rewards and final ratio using the between-subjects factor of Access Condition.

## Results

### Behavioral data

We trained the rats for social self-administration with access to a partner for full contact and screen contact. During training, the rats increased the number of social rewards over time in both conditions (main effect of Session, F _(2.5,45.1)_ = 5.56, p<0.01, Figure S1D), however number of rewards was higher in the full-contact, compared to the screen-contact condition (main effect of Access Condition, F _(1,18)_ = 6.83, p=0.02, Figure S1E). During the progressive-ratio test, the number of social rewards and final ratio were higher in full-contact, compared to the screen-contact condition (main effect of Access Condition, F _(1,18)_ = 5.12, p=0.04, Figure S1F).

### 2. PET data: homecage vs. heroin seeking or social seeking in saline-trained rats

To assess differences in metabolic activity patterns during ‘saline seeking’ and social seeking in saline-trained rats, we compared FDG uptake during homecage scans with uptake during saline- and social-seeking scans. Overall, both saline and social seeking, relative to homecage conditions, were associated with increased FDG uptake across several brain regions. Specifically, both saline and social seeking increased uptake in clusters of voxels in the thalamus (saline seeking: p = 0.041; social seeking: p = 0.013), colliculus (saline seeking: p = 0.087; social seeking: p = 0.021), and cerebellum, with social seeking showing larger and more extensive clusters than saline seeking (Figure S2C-E). Saline seeking was additionally associated with increased uptake in the amygdala (p = 0.039) (Figure S2C-D). Finally, direct comparison between saline- and social-seeking scans revealed increased FDGuptake during social seeking, relative to saline seeking, in clusters including the septum (p = 0.0001), striatum (p<0.001), and ventral subiculum (p<0.001) (Figure S2G-H).

---

**Table S1.** Statistical analysis (SPSS GLM repeated-measures module)

| Figure | Data | Software package | Primary statistic | Main & Interaction effects | F/H/t/r/z statistic | p-value |
| --- | --- | --- | --- | --- | --- | --- |
| **Figure 1C** | Social and Heroin Self-Administration | SPSS | General Linear Model Repeated measure | Social session | $F_{(2.41, 82.1)} = 33.96$ | **p<0.001\*** |
| | | | | Group (Heroin, Saline) | $F_{(1, 34)} = 0.081$ | p=0.777 |
| | | | | Group x Social Session | $F_{(2.41, 82.1)} = 0.36$ | p=0.74 |
| | | | | Heroin session | $F_{(4.71, 160.1)} = 3.05$ | **p=0.013\*** |
| | | | | Group | $F_{(1, 34)} = 9.54$ | **p=0.004\*** |
| | | | | Group x Heroin session | $F_{(4.71, 160.1)} = 28.70$ | **p<0.001\*** |
| **Figure 1D** | Burst Number | SPSS | General Linear Model Univariate | Group (Heroin, Saline) | $F_{(1, 34)} = 30.43$ | **p<0.001\*** |
| **Figure 1E** | Heroin seeking Active lever presses | SPSS | General Linear Model Univariate | Group (Heroin, Saline) | $F_{(1, 34)} = 14.20$ | **p<0.001\*** |
| **Figure 1F** | Social seeking Active lever presses | SPSS | General Linear Model Univariate | Group (Heroin, Saline) | $F_{(1, 34)} = 1.961$ | p=0.170 |
| **Figure 1G** | Preference score during choice sessions | SPSS | General Linear Model Repeated measures | Choice session | $F_{(3.9, 132.6)} = 2.29$ | p=0.065 |
| | | | | Group (Heroin, Saline) | $F_{(1, 34)} = 13.87$ | **p<0.001\*** |
| | | | | Group x Choice session | $F_{(3.9, 132.6)} = 2.09$ | p=0.087 |
| **Figure 1H** | Average Preference score | SPSS | General Linear Model Univariate | Group (Heroin, Saline) | $F_{(1, 34)} = 12.44$ | **p=0.001\*** |
| **Figure 1I** | Heroin seeking Active lever presses | SPSS | General Linear Model Univariate | Group (Heroin, Saline) | $F_{(1, 34)} = 13.06$ | **p<0.001\*** |
| **Figure 1J** | Social seeking Active lever presses | SPSS | General Linear Model Univariate | Group (Heroin, Saline) | $F_{(1, 34)} = 4.78$ | **p=0.036\*** |
| **Figure 2D** | FDG uptake homecage scans in saline vs heroin rats | SPSS | General Linear Model Univariate | Group (Heroin, Saline) - Insula | $F_{(1, 34)} = 7.86$ | **p=0.008\*** |
| | | | | Group (Heroin, Saline) - Somatosensory | $F_{(1, 34)} = 5.56$ | **p=0.024\*** |
| | | | | Group (Heroin, Saline) - Hippocampus | $F_{(1,34)} = 6.16$ | $p=0.018^*$ |
| | | | | Group (Heroin, Saline) - Pons | $F_{(1,34)} = 11.32$ | $p=0.002^*$ |
| **Figure 2F** | FDG uptake heroin seeking scans in saline vs heroin rats | SPSS | General Linear Model Univariate | Group (Heroin, Saline) – Striatum | $F_{(1,34)} = 8.31$ | $p=0.007^*$ |
| | | | | Group (Heroin, Saline) – Septum | $F_{(1,34)} = 7.84$ | $p=0.008^*$ |
| | | | | Group (Heroin, Saline) – Thalamus | $F_{(1,34)} = 10.97$ | $p=0.002^*$ |
| | | | | Group – Cerebellum | $F_{(1,34)} = 16.10$ | $p<0.001^*$ |
| **Figure 2H** | FDG uptake social seeking scans in saline vs heroin rats | SPSS | General Linear Model Univariate | Group (Heroin, Saline) - Retrosplenial | $F_{(1,34)} = 19.24$ | $p<0.001^*$ |
| | | | | Group (Heroin, Saline) – Cerebellum | $F_{(1,34)} = 21.55$ | $p<0.001^*$ |
| **Figure 3D** | FDG uptake homecage vs heroin seeking scans in heroin rats | SPSS | General Linear Model Univariate | Scan Condition (homecage, heroin seek) – Olfactory bulb | $F_{(1,27)} = 118.01$ | $p<0.001^*$ |
| | | | | Scan Condition (homecage, heroin seek) - Hypothalamus | $F_{(1,27)} = 34.47$ | $p<0.001^*$ |
| | | | | Scan Condition (homecage, heroin seek) – Cerebellum | $F_{(1,27)} = 41.31$ | $p<0.001^*$ |
| **Figure 3F** | FDG uptake homecage vs social seeking scans in heroin rats | SPSS | General Linear Model Univariate | Scan Condition (homecage, social seek) – Olfactory bulb | $F_{(1,27)} = 62.24$ | $p<0.001^*$ |
| | | | | Scan Condition (homecage, social seek) - Thalamus | $F_{(1,27)} = 31.18$ | $p<0.001^*$ |
| | | | | Scan Condition (homecage, social seek) – Cerebellum | $F_{(1,27)} = 39.92$ | $p<0.001^*$ |
| **Figure 3H** | FDG uptake heroin vs social seeking scans in heroin rats | SPSS | General Linear Model Repeated measures | Scan Condition (heroin seek, social seek) – Striatum | $F_{(1,27)} = 16.42$ | $p<0.001^*$ |
| | | | | Scan Condition (heroin seek, social seek) - Auditory | $F_{(1,27)} = 11.02$ | $p=0.003^*$ |
| **Figure 4C** | Severity vs Piriform Cortex Uptake | SPSS | Pearson Correlation | Across groups | R=-0.39 | **p=0.038\*** |
| **Figure 4D** | Severity vs VHipp Uptake | SPSS | Pearson Correlation | Across groups | R=0.55 | **p=0.002\*** |
| **Figure 4F** | Severity vs Post Sub Uptake | SPSS | Pearson Correlation | Across groups | R=-0.65 | **p<0.001\*** |
| **Figure 4G** | Severity vs Cerebellum Uptake | SPSS | Pearson Correlation | Across groups | R=-0.53 | **p=0.003\*** |
| **Figure 4I** | Severity vs VisualCx Uptake | SPSS | Pearson Correlation | Across groups | R=-0.49 | **p=0.008\*** |
| **Figure 5E** | FDG uptake homecage scans in high and low addicted rats | SPSS | General Linear Model Univariate | Group (high, low) – PirCx | F (1,13) = 16.25 | **p=0.001\*** |
|  |  |  |  | Group (high, low) - NAc | F (1,13) = 7.53 | **p=0.017\*** |
|  |  |  |  | Group (high, low) - LHA | F (1,13) = 18.79 | **p<0.001\*** |
|  |  |  |  | Group (high, low) - vHipp | F (1,13) = 20.29 | **p<0.001\*** |
| **Figure 5G** | FDG uptake heroin seeking scans in high and low addicted rats | SPSS | General Linear Model Univariate | Group (high, low) – Postsub | F (1,13) = 5.96 | **p=0.03\*** |
|  |  |  |  | Group (high, low) - Cerebellum | F (1,13) = 15.94 | **p=0.002\*** |
| **Figure S1D** | Social and Heroin Self-Administration | SPSS | General Linear Model Repeated measure | Social session | F (2.5,45.1) = 5.56 | **p=0.004\*** |
|  |  |  |  | Social Condition (Partial, Full Contact) | F (1,18) = 6.83 | **p=0.018\*** |
|  |  |  |  | Social Condition x Social Session | F (2.5,45.1) = 1.53 | p=0.23 |
| **Figure S1E** | Average Rewards | SPSS | General Linear Model Univariate | Social Condition (Partial, Full Contact) | F (1,18) = 6.83 | **p=0.018\*** |
| **Figure S1F** | PR active lever presses and final ratio | SPSS | General Linear Model Univariate | Social Condition (Active Lever Presses) | F (1,18) = 5.105 | **p=0.037\*** |
|  |  |  |  | Social Condition (Final Ratio) | F (1,18) = 5.19 | **p=0.035\*** |
| **Figure S2D** | FDG uptake homecage vs saline seeking scans in saline rats | SPSS | General Linear Model Repeated measure | Scan Condition (homecage, heroin seek) – Amygdala | F (1,7) = 6.39 | **p=0.039\*** |
|  |  |  |  | Scan Condition (homecage, heroin seek) - Thalamus | F (1,7) = 6.25 | **p=0.041\*** |
| | | | | Scan Condition (homecage, heroin seek) – Colliculus | $F_{(1,7)} = 3.97$ | $p=0.087$ |
| **Figure S2F** | FDG uptake homecage vs social seeking scans in saline rats | SPSS | General Linear Model Repeated measure | Scan Condition (homecage, social seek) – Thalamus | $F_{(1,7)} = 10.80$ | **$p=0.013^*$** |
| | | | | Scan Condition (homecage, social seek)- Colliculus | $F_{(1,7)} = 8.75$ | **$p=0.021^*$** |
| | | | | Scan Condition (homecage, social seek) – Cerebellum | $F_{(1,7)} = 13.62$ | **$p=0.008^*$** |
| **Figure S2H** | FDG uptake homecage vs social seeking scans in saline rats | SPSS | General Linear Model Repeated measure | Scan Condition (heroin seek, social seek) – Septum | $F_{(1,7)} = 25.98$ | **$p=0.001^*$** |
| | | | | Scan Condition (heroin seek, social seek) - Striatum | $F_{(1,7)} = 34.18$ | **$p<0.001^*$** |
| | | | | Scan Condition (heroin seek, social seek) – vSub | $F_{(1,7)} = 66.31$ | **$p<0.001^*$** |

## Data availability statement

The data that support the findings of this study are available from the corresponding authors upon reasonable request.

## Acknowledgment statement

This research was supported [in part] by the Intramural Research Program of the National Institutes of Health(NIH). The contributions of the NIH author(s) are considered Works of the United States Government. The findings and conclusions presented in this paper are those of the author(s) and do not necessarily reflect the views of the NIH or the U.S.Department of Health and Human Services

## Author contributions

All authors contributed to different aspects of the study, including the design and performance of the research, data analysis, and write-up of the paper.

## Funding

The research was supported by funds from the Intramural Research Program of the NIDA-NIH (YS, grant number, 1ZIADA000434-25; MM, grant number, ZIADA000069; TK, grant number, ZIADA000642). The funding body did not play a role in the design of the study and collection, analysis, and interpretation of data, and in writing the manuscript.

## Conflict of interest disclosure

The authors declare that they do not have any conflicts of interest (financial or otherwise) related to the content of the paper.

## Competing interests’ statement

The authors declare no competing financial interests.

## Ethics approval statement

Procedures were approved by local Ethical Committee of the Intramural Research Program of NIDA, NIH.

## Notes

### Competing Interest Statement

The authors have declared no competing interest.

